# Single-cell analysis of Non-CG methylation dynamics and gene expression in human oocyte maturation

**DOI:** 10.1101/651141

**Authors:** Bo Yu, Naresh Doni Jayavelu, Stephanie L. Battle, Thomas H. Smith, Samuel E Zimmerman, Timothy Schimmel, Jacques Cohen, Jessica C. Mar, R. David Hawkins

**Affiliations:** Department of OBGYN, University of Washington School of Medicine, Seattle, WA 98195, USA; Departments of Medicine and Genome Sciences, University of Washington School of Medicine, Seattle, WA 98195, USA; Department of Systems and Computational Biology, Albert Einstein College of Medicine, Bronx, NY 10461, USA; Reprogenetics LLC, Livingston, NJ 07039, USA

**Keywords:** DNA methylation, oocyte maturation, gene expression

## Abstract

Oocyte maturation is a coordinated process that is tightly linked to reproductive potential. A better understanding of gene regulation during human oocyte maturation will not only answer an important question in biology, but also facilitate the development of *in vitro* maturation technology as a fertility treatment. We generated single-cell transcriptome and use previously published single-cell methylome data from human oocytes at different maturation stages to investigate how genes are regulated during oocyte maturation, focusing on the potential regulatory role of non-CG methylation. *DNMT3B*, a gene encoding a key non-CG methylation enzyme, is one of the 1000 genes upregulated in mature oocytes, which may be at least partially responsible for the increased non-CG methylation as oocytes mature. Non-CG differentially methylated regions (DMRs) between mature and immature oocytes have multiple binding motifs for transcription factors, some of which bind with *DNMT3B* and may be important regulators of oocyte maturation through non-CG methylation. Over 98% of non-CG DMRs locate in transposable elements, and these DMRs are correlated with expression changes of the nearby genes. Taken together, this data indicates that global non-CG hypermethylation during oocyte maturation may play an active role in gene expression regulation, potentially through the interaction with transcription factors.

## INTRODUCTION

Proper development of the mature oocyte is an essential part of reproduction and the prerequisite for fertilization and downstream embryonic development. In humans, oocytes are arrested in prophase of meiosis I and remain quiescent until decades later when the follicles are recruited for growth^1^. Oocytes must mature and undergo transcription and physiological changes in preparation for ovulation and fertilization. When the arrested prophase I oocyte (also known as germinal vesicle stage) is recruited to grow and mature, the germinal vesicle breaks down and the oocyte resumes meiosis I, leading to the intermediate maturation state metaphase I (MI stage). As meiosis II progresses, the oocyte divides into a polar body and a mature metaphase II oocyte (MII stage). The MII oocyte is ready for ovulation and fertilization, and carries the maternal genome into the embryo after fertilization. The gene regulatory dynamics associated with oocytes maturation are of great interest to developmental biology and for clinical development of better *in vitro* maturation technology as a fertility treatment.

The transcriptomic landscape of oocyte development has been described in several mammalian species including cow, rabbit, rhesus, mouse and human^2, 3^. Most of these studies, however either used microarray data or focus on only 2 stages of oocyte maturation, the germinal vesicle (GV) and the metaphase II oocyte (MII)^4^. From these studies we have learned that there are fewer transcripts in the MII oocyte which can be due to both reduced expression and RNA degradation^2, 5^. Thousands of differentially expressed genes, both upregulated and downregulated, have been identified from GV to MII^2, 5, 6^. Further elucidation of gene regulatory mechanisms would help us understand the transcriptomic dynamics of oocyte maturation.

Similarly, previous DNA methylation studies on developing oocytes have provided great insight to the epigenomic landscape of the maturing oocyte. Mouse studies have shown that there is an overall increase in methylation as oocytes grow and mature^7, 8^. In humans, the process of DNA methylome erasure in primordial germ cells and in pre-implantation embryos was recently shown in several genome-wide studies^9, 10^. Our group published data in single-cell DNA methylome of human oocytes at various maturation stages last year, and demonstrated genome-wide increase in non-CG methylation as oocytes mature. Few previous study have investigated the epigenomic regulatory mechanisms that control differential gene expression during human oocyte maturation from GV to MII. The biological role of genome-wide non-CG methylation remodeling in the final stage of oocyte maturation remains to be investigated.

Most previously published human oocyte gene expression data either looks at only two timepoints, focuses their comparisons on MII versus *in vitro* matured oocytes, or focuses their comparison on oocyte versus early embryo^2, 6, 11–13^. Here we generated and correlated single-cell mRNA-seq and single-cell whole genome bisulfite sequencing data from the same individual cohort at three oocyte maturation stages (GV, MI, MII), in order to gain a better understanding gene regulation during human oocyte maturation through DNA methylation. We find that the accumulation of non-CG methylation in mature MII oocytes is accompanied by upregulation of the *DNMT3B* gene. There was a correlation between differentially methylated regions (DMRs) and gene expression with distal DMRs negatively correlating with gene expression and gene body DMRs positively correlating with gene expression. We identify transcription factors (TFs), ETS1 and YY1, which have binding motifs at DMRs and potentially direct DNMT3B methylation in the maturing oocyte genome. Many of the DMRs are located in the regulatory regions of the genome. These results suggest a regulatory role of non-CG methylation in transcriptomic changes during oocyte maturation.

## RESULTS

### Gene Expression, DEGs and Pathways

Oocytes were collected from 17 donors for single-cell transcriptome and single-cell DNA methylome (Table S1). Some of the oocytes were obtained from the same individuals. In total, 21 mRNA-seq libraries were generated and analyzed together with previously published data on 32 whole genome bisulfite sequencing libraries^14^. For transcriptome data processing, mRNA sequencing reads were mapped to hg19 and Cufflinks^15^ was used to calculate FPKM values. Individual samples were assessed for quality and uniformity based on their distribution of FPKM values (Figure S1A and S1B). We found high correlation between FPKM and normalized read count (Figure S1C) and therefore decided to continue all downstream analysis with FPKM values. We calculated the pairwise Pearson’s correlation coefficient to assess intra-versus inter-individual variation (Figure S1D). Since we did not observe higher correlations between samples from the same individual compared to across individuals at the same oocyte maturation stage, we merged samples of the same stage and used the merged FPKM values for all further analyses. We asked how similar the oocyte maturation stages were based on their transcriptome. Principal component analysis of the top 1000 expressed genes shows a clear separation between the immature GV/MI oocytes and the mature MII oocytes, with the GV and MI stages being almost indistinguishable from each other (Figure 1A). We next looked at how many expressed genes could be found in each maturation stage. We observed 15,224 GV, 13,283 MI and 10,892 MII expressed genes (FPKM >= 1; Figure 1B). We asked how many genes were specific to each stage and found 427 GV, 3 MI and 7 MII specific genes. Such few cell-type specific genes in the later stages suggests that *de novo* transcription occurs in the earliest stage of oocyte maturation^16, 17^. All 4 zona pellucida glycoprotein genes (*ZP1*, *ZP2*, *ZP3*, and *ZP4*) were highly expressed in all three stages. We were also able to validate oocyte specific genes *DAZL*, *GDF9*, and *BMP15* as being expressed and *RBBP7* as significantly highly expressed in MII oocytes compared to MI (log 2 fold change 1.44; Figure S1E). Some of the highest expressed genes in MII oocytes were *PTTG1* and *TUBB8*. *PTTG1* encodes a securin protein that prevents sister chromatid separation. *TUBB8* encodes the beta-tubulin subunit primarily expressed in oocytes and the early embryo. Mutations in *TUBB8* have been identified in infertile women with oocyte maturation arrest^18–20^. These genes have clear relevance and significance in oocyte maturation and development.

**Figure 1:**
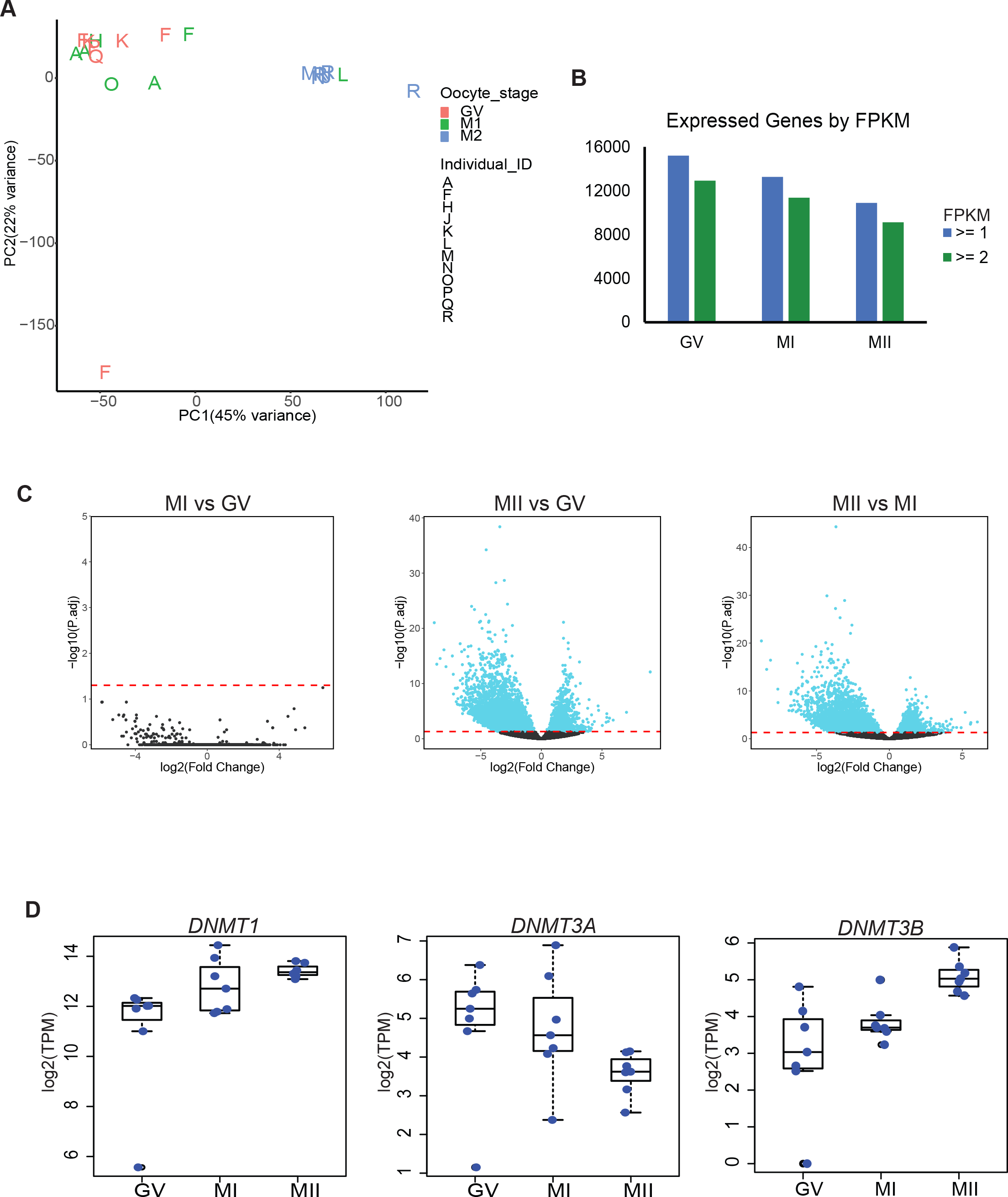
Gene expression in GV, MI and MII oocytes. A. PCA of top 1000 expressed transcripts in individual RNA-Seq libraries. Letters on plot correspond to individual sample IDs and color corresponds to oocyte stage. B. Bar chart of total number of transcripts expressed at equal to or less than 1 or 2 FPKM cutoffs from merged single-cell RNA-Seq datasets. C. Volcano plot of differential gene expression in pairwise comparisons. Dashed line is adjusted p-value cutoff of 0.05. D. Bar chart of FPKM values of DNMTs. Each blue dot is the FPKM value of each individual sample.

We used DESeq2^21^ to calculate the number of differentially expressed genes (DEGs) between the three oocyte maturation stages. The largest number of DEGs was observed between the immature stages and MII (Figure 1C). There were 5,979 DEGs between GV and MII and 4,835 between MI and MII with an adjusted p-value of equal to or less than 0.05. There were no significant DEGs between GV and MI, supporting the results of our PCA, that these two oocyte stages are very similar at the level of protein coding genes. This was also previously observed in microarray data on human oocytes^6, 17, 22^. In order to ensure our findings were not due to a bias in our analysis, we repeated the DEG analysis using a different program, edgeR^23^. Even though edgeR called fewer DEGs, there was a high overlap with those called by DESeq2 (Figure S1F).

Of the DEGs we identified using DESeq2, 1,362 were upregulated in MII compared to GV and 4,617 were downregulated. In the MII-to-MI pairwise comparison, 1,077 were upregulated and 3,758 were downregulated. We focused on the MII-to-MI DEGs as these two timepoints mark the transition from immature to mature oocyte (Table S2). The upregulated genes fell into pathways involving RNA degradation, splicing and transport (Table 1). Cell cycle, ubiquitin mediated proteolysis and oocyte meiosis were also significantly enriched pathways. These pathways are anticipated since genes involved in regulating meiosis should be expressed in the MII oocyte. Genes involved in pathways involving RNA dynamics are necessary given that: (1) the oocyte accumulates maternal RNAs necessary for early embryonic development, (2) there is slow maternal RNA degradation during oocyte maturation, as detected by the reduction in transcripts, and (3) maternal RNA degradation occurs shortly after fertilization. Downregulated pathways included various metabolic pathways including the TCA cycle and oxidative phosphorylation. We speculate that the downregulation of TCA and oxidative phosphorylation genes could be influenced by the underdeveloped mitochondria in oocytes^24, 25^.

**Table 1.** Listed are significantly enriched KEGG pathways for up-and downregulated DEGs (differentially expressed genes) in the MII to MI pairwise comparison.

MII vs MI DEG KEGG Pathways
| MII Upregulated DEGs |  |
| --- | --- |
| Pathway | Adj p-value |
| Ribosome | 1.85E-07 |
| RNA degradation | 0.0000112 |
| Spliceosome | 0.0001546 |
| Cell cycle | 0.0001546 |
| RNA transport | 0.00652 |
| Ubiquitin mediated proteolysis | 0.01526 |
| Oocyte meiosis | 0.03 |
| MII Downregulated DEGs |  |
| Pathway | Adj p-value |
| Metabolic pathways | 6.98E-08 |
| Oxidative phosphorylation | 5.72E-07 |
| Parkinson's disease | 0.0002435 |
| Carbon metabolism | 0.001298 |
| 2-Oxocarboxylic acid metabolism | 0.001929 |
| Huntington's disease | 0.003961 |
| Pyrimidine metabolism | 0.004119 |
| Non-alcoholic fatty liver disease | 0.004944 |
| Proteasome | 0.004944 |
| Biosynthesis of amino acids | 0.006568 |
| SNARE interactions in vesicular transport | 0.01449 |
| Alzheimer's disease | 0.01509 |
| Glyoxylate and dicarboxylate metabolism | 0.01581 |

Of the genes most upregulated in mature MII oocytes we found *SLX1B* and *MATN3* to be among the highest upregulated protein coding genes (log2 fold change 5.2 and 4.6 respectively). *SLX1B* encodes a protein that resolves DNA secondary structures that result from DNA repair or recombination, which we propose would be necessary for developing from MI to MII and completing meiosis post-fertilization. The MATN3 protein is a component of extracellular matrix which we hypothesize may be important in the development of oocyte extracellular structures. We queried our RNA-seq data for expression level of epigenetic regulators (Figure S1G and Figure 1D). *TET3* is significantly upregulated in MII oocytes with a log2 fold change of 3.3x compared to MI oocytes. *TET3* and *TET2*, which are expressed in all 3 stages of oocyte maturation, are important in removing methylation in the zygotic genome upon fertilization^26, 27^. *DNMT1* is highly expressed in all 3 oocyte maturation stages and *DNMT3B* is significantly upregulated 2.8x in MII compared to MI oocytes. *DNMT3A* expression is reduced in MII and there is no detectable *DNMT3L*. The upregulation of *DNMT3B* led us to take a closer look at DNA methylation trends in the maturing oocyte.

### DNA Methylation Correlates with Gene Expression in Oocytes

Global methylation trends for this cohort were previously described^14^. However, with the addition of transcriptomic data, we can investigate associations between DNA methylation and gene expression in oocytes. GV and MI stage oocytes had similar average methylation levels for cytosines in all contexts, CHG, CHH and CpG. We observed similar average CpG methylation levels across all three stages (Figure S2A).

The non-CG average methylation level doubled in MII compared to the immature stages. The increase of non-CG methylation only in the MII stage coincides with the upregulation of *DNMT3B* in MII. Gene body methylation has been shown to be positively correlated with gene expression{Lister, 2009}. We asked if this same phenomenon can be observed in all three oocyte growth stages. There is a minor, but positive, correlation between gene body methylation and gene expression for all C contexts in all three stages (Figure 2A and Figure S2B). This correlation is stronger for methylated CpGs, which have higher average methylation levels than non-CGs.

**Figure 2:**
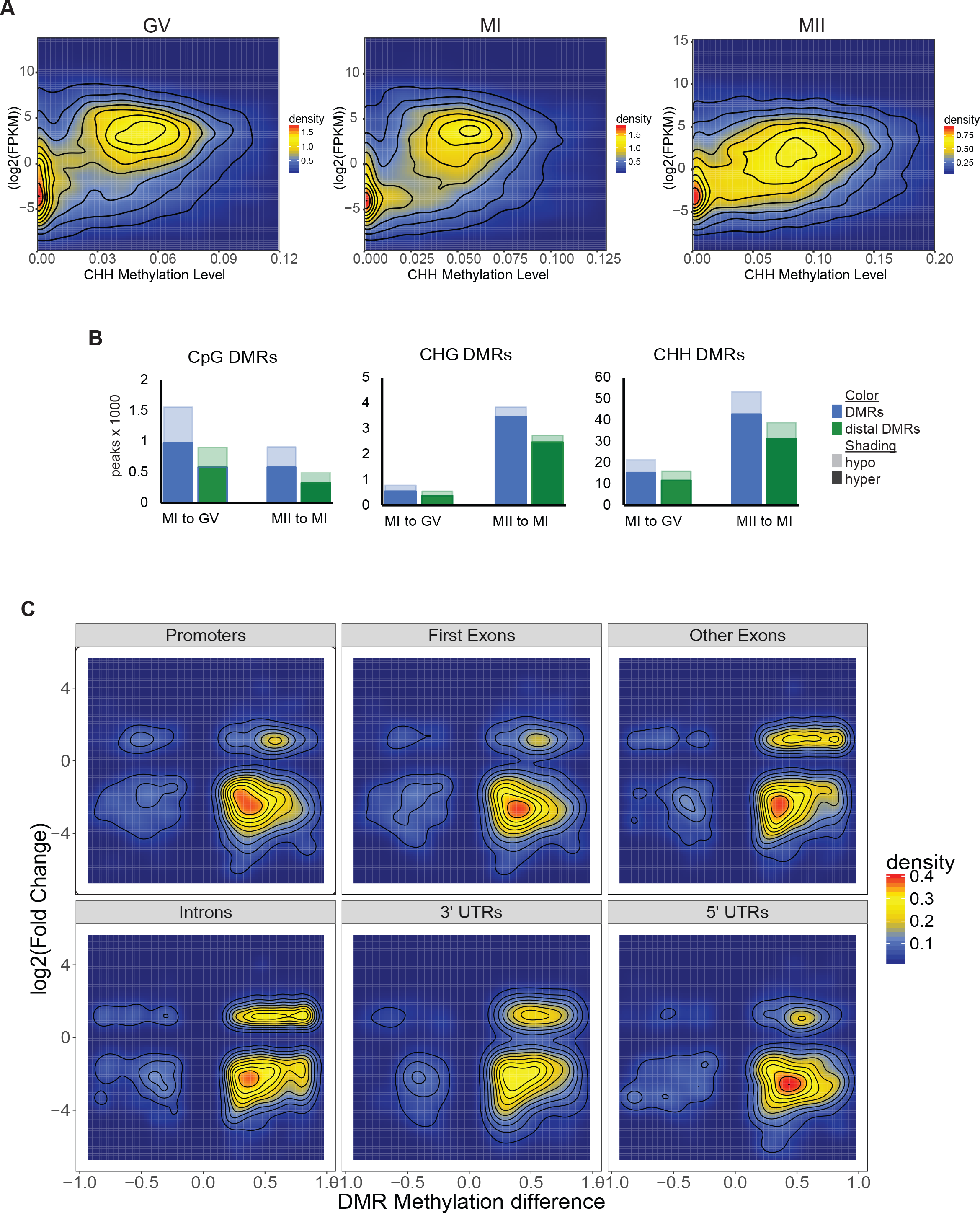
Gene body methylation and Expression Correlations. A. Density plots of gene body methylation levels for CHH context and the corresponding gene’s expression level in GV, MI and MII oocytes. Density color scale to the right of each respective plot. B. Differentially methylated region (DMR) counts by the thousands for each C context in GV, MI and MII oocytes. Blue bar to the left is total DMR count. Green bar to the right is the count of distal DMRs. Hypermethylated DMRs are solid color while hypomethylated DMRs are lightly shaded. C. Methylation level of MII-to-MI CHH DMRs in specific genic regions and their corresponding gene expression level. Scale for density plot is to the right.

### Differentially Methylated Regions Correlate with Gene Expression in Oocyte Maturation

In order to further investigate methylation changes during oocyte maturation, we identified differentially methylated regions (DMRs) for each C context. We took differentially methylated regions from Yu et al., 2017, merged overlapping regions and used those for all downstream analysis. Only DMRs from the same stage oocyte were merged. DMRs have been proposed to be *cis*-regulatory elements in other cell types^28^, therefore we asked if the same could be true in the maturing oocyte. We counted over 70% of non-CG DMRs as distal, meaning not overlapping a gene promoter (−2kb to +500bp of gene TSS), exons or UTRs. All distal DMRs also had over 80% overlap with ENCODE DNaseI hypersensitivity sites (Table S3). The majority of DMRs were hypermethylated in the mature oocyte in the pairwise comparisons to the immature stages (Figure 2B and Table S3). Hypermethylated DMRs in all genic context were associated with downregulation of gene expression in the MII-to-MI comparison (Figure 2C and Figure S2C). However some hypermethylated CHH DMRs, specifically in introns and other exons, are associated with upregulated gene expression. In fact, 603 (60%) of the upregulated MII-to-MI DEGs have a DMR in their gene body. Further suggesting a role of DMRs in gene regulation at this stage of development.

### DNMT3B binding partners and hypermethylation of gene targets

Of the DNMTs, only *DNMT3B* is upregulated as oocytes mature. DNMT3B is one of the *de novo* methyltransferases and can methylate non-CG motifs^29^. DNA methylation as a mechanism of gene regulation has been intensively studied in embryonic, germ and somatic cells. The upregulation of *DNMT3B* and increase of non-CG methylation could be necessary for several mechanisms in the developing oocyte. One potential mechanism could be to methylate the maternal genome in preparation for fertilization and embryo development. The second could be to regulate expression of oocyte genes during maturation. Many studies have examined this first point and looked at the epigenome from the MII oocyte through the beginning of embryogenesis^9, 30^. We chose to focus on the second point.

In order to gain a better understanding of what factors may direct DNMT3B to methylate regions in the maturing oocyte genome, we looked for transcription factors (TF) that have been previously shown to directly bind DNMT3B via protein-protein binding array^31^ (Figure 3A). Using the protein-protein binding data from Hervouet et al. 2009, we filtered for TFs that are expressed in oocytes (Figure 3B). Of the DNMT3B-TF binding partners 29 were expressed in MI or MII stage oocytes. Expression of three of DNMT3B-TF binding partners were significantly upregulated in MII compared to MI: *ATF2*, *CREB1* and *SP4*. We next looked for transcription factor binding motifs in MII-to-MI distal DMRs (Table S4). We further filtered our list of expressed DNMT3B binding partners for factors that have their known motif in an MII distal DMR (denoted with * in Figure 3B) and of those only ETS1 and YY1 were expressed in MII.

**Figure 3:**
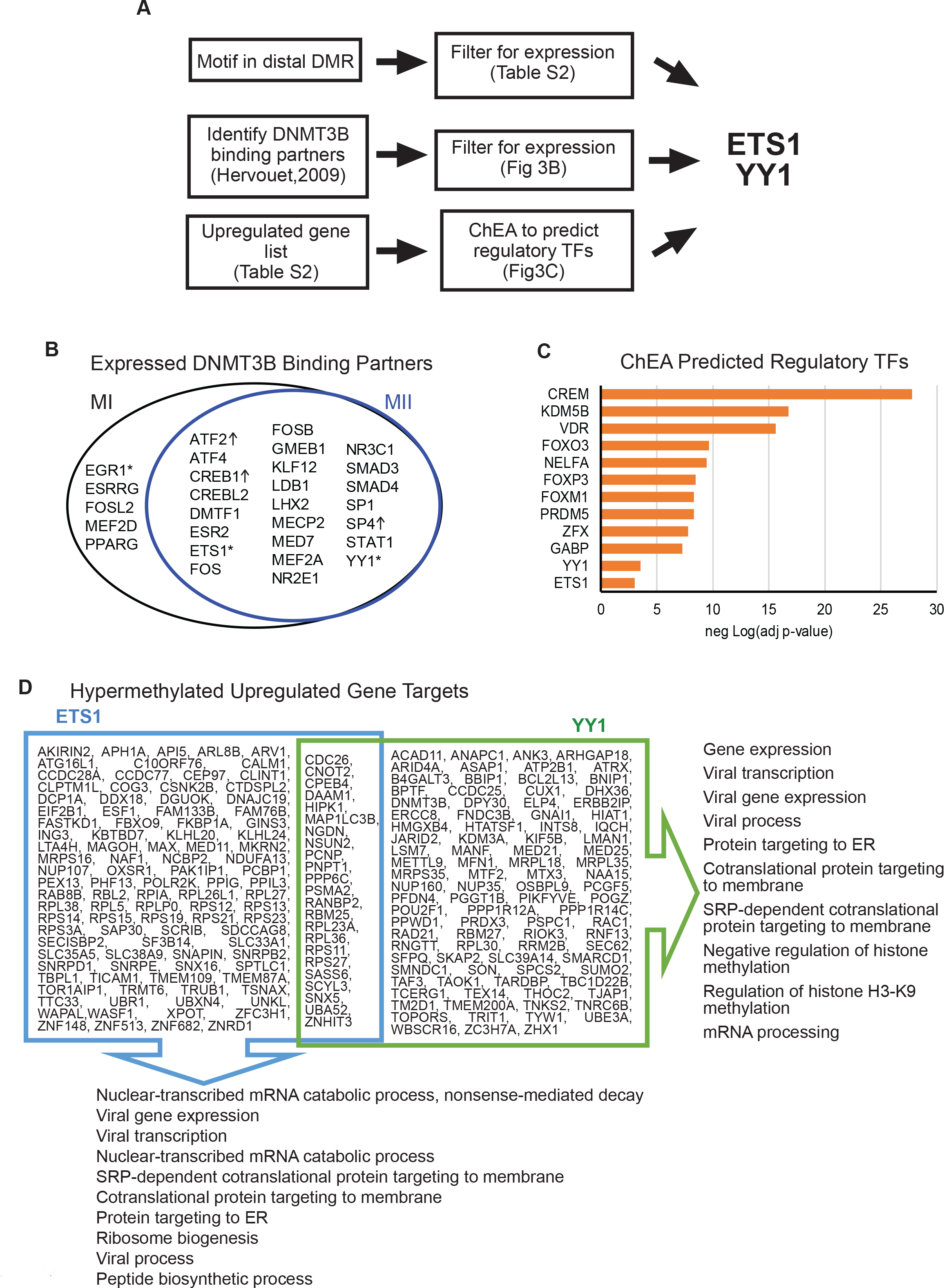
Regulatory Transcription factors at differentially methylated regions and their gene targets. A. Flow chart of the two different approaches that identified ETS1 and YY1 as regulatory transcription factors (TF) in oocytes. B. Venn diagram showing DNMT3B binding partners that are expressed in MI or MII (FPKM >= 1). Of the expressed transcription factors (TFs), only three (denoted with “*”) have their binding motifs present in MII-to-MI DMRs. TF with “↑” are significantly upregulated in MII compared to MI. C. Bar charts of adjusted p-value of select TFs predicted to regulate MII-to-MI downregulated genes by ChEA. Orange bars have adjusted p-value less than 0.05. D. MII-to-MI downregulated genes predicted to be regulated by ETS1 and YY1. The top ten GO Biological Processes for each gene list are displayed.

In a complementary approach, we ran our list of upregulated genes through ChIP Enrichment Analysis (ChEA)^32^ which predicts regulatory TFs from a gene list (Figure 3A). ETS1 and YY1 were predicted to be regulatory TFs by ChEA with statistically significant enrichment scores (adjusted p-value 9.0*10^-4^ and 2.9*10^-4^; Figure 3C). Notably, we also found that ETS1 (ChEA adjusted p-value 5.3*10^-17^) and YY1 (ENCODE TF ChIP-seq adjusted p-value 2.4*10^-21^) were predicted to be regulatory TFs in the downregulated gene list as well. Therefore, we took a closer look at ETS1 and YY1 and their regulatory role in MII stage oocytes.

ETS1 is a transcription factor involved in many cellular processes including tumorigenesis and stem cell development. ETS family proteins are phosphorylated by MAP kinases and their activity as either an activator or repressor is modulated by co-factor binding^33^. YY1 is a ubiquitous transcription factor shown to have both activator and repressor activity^34, 35^. YY1 can interact with other proteins to stabilize enhancer-promoter loops^35, 36^. More importantly, YY1 is required for oocyte growth and follicular expansion^37^. Mice with a conditional knockout of YY1 are infertile and lack mature oocytes^37^. Using ChEA data, we identified the upregulated gene targets of ETS1 and YY1 (Figure 3D). There were 123 gene targets of ETS1 and 117 gene targets of YY1 that were upregulated in MII compared to MI. Common GO Biological Processes between the two gene lists included mRNA processes, viral gene expression and protein targeting particularly to the ER. YY1 regulated genes were also involved in epigenetic regulation, specifically of histone H3 K9 methylation. Of the 603 MII upregulated genes that have a DMR in their gene body, 62 genes had an ETS1 binding site in their DMR and 151 genes had an YY1 binding site, with 60 genes in common between these two groups. Additionally, we found these gene body DMRs to be hypermethylated in 58 of the genes with ETS1 and 140 of the genes with YY1 binding sites. Therefore the gene body hypermethylation could be a result of ETS1 and YY1 directing DNMT3B to those genic regions.

### LINE1 Elements at DMRs

Transposable elements (TEs) have been shown to function as regulatory elements and are expressed in oocyte^9, 38–40^. Since TEs have been shown to be hypermethylated in MII oocytes compared to pre-implantation embryo^9^, we asked if there was differential methylation at TEs during oocyte maturation. We used TE coordinates from RepeatMasker to assess how many DMRs were at TE elements in the genome. Over 95% of DMRs, and over 98% of non-CG DMRs, were at transposable elements in the genome (Table S3). We focused further of LINE1 elements as those have been shown to be expressed in mammalian oocytes and early embryogenesis^40, 41^. We used GREAT to predict genes that were potentially regulated by the LINE1 elements in MII non-CG hypermethylated distal DMRs (Figure S3). A number of terms were driven by the proximity of LINE1 elements to the *AR* gene. When we looked in the UCSC genome browser, there are cluster of non-CG hypermethylated LINE1 elements upstream of the *AR* gene (Figure 4). In our gene expression data *AR* is expressed in GV and expression drops until it is undetectable in MII oocytes. *AR* expression is important in early follicle maturation and in knockout mouse models, females have many phenotypes associated with reduced fertility^42^. AR is also important for germinal vesicle breakdown, the process by which the GV oocyte resumes meiosis to become the mature MII oocyte^42^. *TMEFF2* is also a gene potentially regulated by proximal hypermethylated LINE1 elements (Figure 4). *TMEFF2* is upregulated in early oocyte development in the primordial/primary follicle stage^43^ and in our data we observed downregulation of TMEFF2 from GV to MII. Therefore non-CG hypermethylation in the MII stage oocyte could be a means to suppress LINE1 elements that help regulate the expression of early oocyte maturation genes.

**Figure 4:**
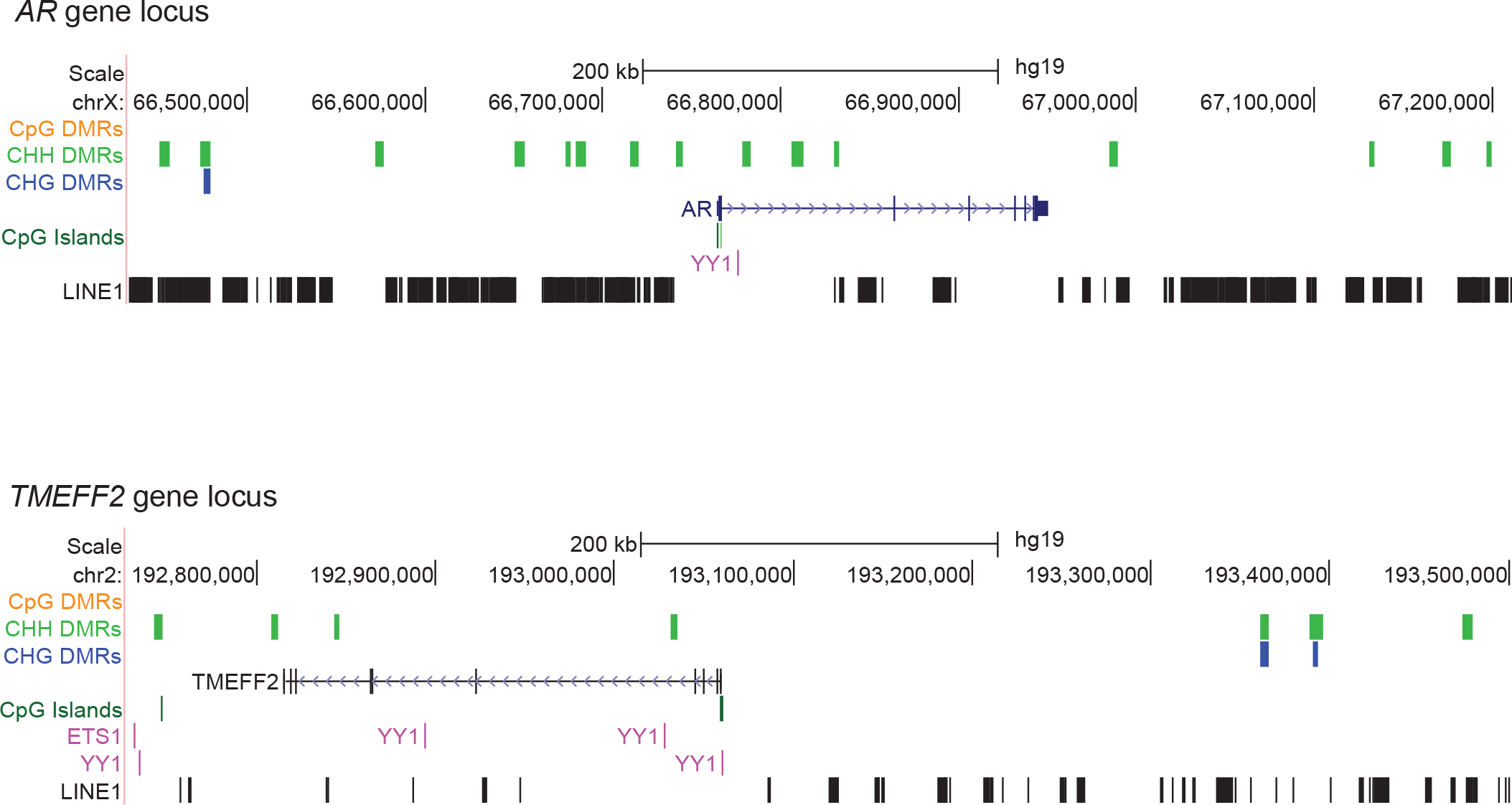
Genes Regulated by Hypermethylated non-CG DMRs at LINE1 elements. UCSC browser images of the *AR* and *TMEFF2* loci. Hypermethylated DMRs for each context are shown and the location of LINE1 elements are also displayed.

## DISCUSSION

While many studies have looked at the genetic and epigenetic contributions of the oocyte to embryonic development, less focus has been on regulatory factors during oocyte maturation. Here we used RNA-seq and WGBS data generated in single-cell oocytes at three different timepoints of oocyte maturation. We use oocytes from patients undergoing ART and therefore one major limitation of our study is the potential heterogeneity of the patients samples. In our three timepoints we observed less RNA transcripts and general downregulation of gene expression as the oocyte matures. However there are genes that are specifically upregulated in mature MII oocytes and these include epigenetic regulators. The upregulation of *DNMT3B* is accompanied by the increase in non-CG methylation in the mature MII genome. Most of the MII-to-MI DMRS, particularly non-CG DMRs, exist in the regulatory region of the genome and may be involved in regulating expression of oocyte maturation genes, partially through suppression of LINE1 elements. We observe several TF motifs in the MII/MI DMRs of which, the transcriptional regulators ETS1 and YY1 have previously been shown to interact with DNMT3B based on protein array binding data from Hervouet et al., 2009. Upregulated gene targets of ETS1 and YY1 are involved in mRNA processes, viral gene expression and protein targeting. We hypothesize that one function of increased non-CG methylation in the maturing oocyte is to regulate genes using TF-DNMT3B directed methylation at MII DMRs. We acknowledge however that testing these regions for regulatory activity is conceivable but nevertheless challenging in this system.

In oocyte development, the increase of non-CG methylation is specific to maturing oocytes. In mice, newborn non-growing oocytes are mostly depleted of non-CG methylation but gain it as cell enter the GV stage^8^. The establishment of non-CG methylation in the immature stages is dependent on Dnmt3a and Dnmt3L activity, while loss of Dnmt3b has no effect^8^. While we do not detect expression of DNMT3L in human oocytes, DNMT3A is upregulated in the immature stages. There is a switch from DNMT3A to DNMT3B expression once oocytes enter the mature MII stage. This is accompanied by an increase in non-CG methylation. We speculate that in human DNMT3B activity may be specific to the mature oocyte.

Intragenic gene regulation via DNMT3B has been observed in other cell-types. For example mutations in DNMT3B occur in the majority of patients with Immunodeficiency, Centromere instability and Facial anomalies (ICF) syndrome. In these patients intragenic binding of DNMT3B was shown to affect expression of transcript isoforms{Gatto,2017}. Gene repression via DNMT3B-TF directed methylation has been observed in other cell types. Dnmt3B interacts with the repressor E2F6 to methylate promoters and silence germline genes in murine somatic tissues^44^. DNMT3B can also interact with PU.1 to methylate promoters of genes during monocyte to osteoclast differentiation^45^. DNMT3B recruitment to genomic loci may also alter the chromatin structure as DNMT3B has been shown to co-IP with HDAC1 and HDAC2 in human and mouse cell lines^46^. One of the TFs we identified in our study YY1, is essential for oocyte maturation and is known act as either an activator or repressor thru interaction with other regulatory proteins ^35, 37^. Here we propose a model where DNMT3B recruitment by TFs, such as ETS1 and YY1, is responsible for targeted gene regulation in the MII oocyte.

Transposable element activity in germ cell and early embryonic development are widely studied^41, 47^. Previously it has been shown that LINE1 activity in GV oocytes is essential for progression to the MII stage^41^. Here we shown a clear example of how LINE1 elements may function as regulatory elements during oocyte maturation. LINE1 elements have previously been shown to be hypermethylated in gamete^9^ thus this hypermethylation must occur during oocyte maturation from GV to MII. Overall this is an example of how hypermethylation of regulatory elements can be used to control gene expression in the maturing oocyte.

Understanding the gene regulatory network in maturing oocytes could provide insight into the etiology of syndromes like oocyte maturation arrest^48^. By generating next generation sequencing data in three maturation timepoints we are providing a resource to the scientific community for further investigation into oocyte development.

## MATERIALS AND METHODS

### Oocyte collection

All human oocytes used in this study were obtained in embryology laboratories at Saint Barnabas Center for Reproductive Medicine and Sher Institutes for Reproductive Medicine under the regulatory oversight of Institutional Review Board (IRB)-approved Human Subjects protocol at each institution. After oocyte retrieval procedures under standard Assisted Reproductive Technology protocols, oocytes that were destined to be discarded were collected under previously obtained written informed consent. All consented materials were donated anonymously and carried no personal identifiers. None of the oocytes was exposed to sperms or discarded due to fertilization or quality issues. GV and MI oocytes were collected at the time of maturity check. Most of the MII oocytes were collected from oocyte donors who had excess oocytes to dispose, due to rare circumstances that removed them from the donor list.

To eliminate contamination from cumulus cells and other cell types, we removed zona pellucida from each oocyte using either acid Tyrode’s solution or mechanical separation techniques. Each oocyte was washed in PBS 2-3 times and immediately frozen in 2 μL of PBS in a −80°C freezer until shipment on dry ice. Oocytes were received for further studies at Albert Einstein College of Medicine (AECOM) under the approval of AECOM IRB, which deemed the project exempt under 45 CRF 46.102(f). All oocytes were processed by the same embryologist (T.S.), including the removal of zona pellucida, washing, cryopreserving, and shipping. All single-cell WGBS and mRNA-seq experiments were carried out by the same individual (B.Y.).

### Library preparation and mRNA-seq

From each single oocyte, cDNA was synthesized using SMART-Seq v4 ultra low input RNA kit (Clontech, cat # 634889). An mRNA-seq library was then prepared from each sample using standard Illumina TruSeq protocol. All libraries were multiplexed and sequenced on the same flow cell in order to minimize sequencing batch effect. Sequencing was performed at the Epigenomics and Genomics Shared Facility at Albert Einstein College of Medicine.

### Gene expression data analysis

The sequence data was aligned to the human genome build 19 using the STAR aligner with default parameters. Gene expression values were computed in FPKM units using Cufflinks^15^. Differentially expressed genes were calculated using DESeq2^21^ and edgeR^23, 49^. Only DEseq2 calculated differentially expressed genes were used for further analysis.

Gene lists were put into Enrichr^50, 51^ to obtain Gene Ontology terms, KEGG pathways, ChEA and ENCODE Transcription factor ChIP-seq enrichment scores.

### WGBS and differentially methylated regions analysis

WGBS data was taken from Yu et al., 2017, where methods and equations are described in depth. Briefly, raw sequence reads were trimmed for adapter contamination using Trim Galore! and mapped to hg19 using Bismark. Lambda phage DNA spike-ins were used to calculate bisulfite conversion efficiency. All conversion efficiencies were above 98%. Methylation levels were called using Bismark Methylation Extractor and methylation levels were calculated for each context (CpG, CHG, CHH) separately. Differential methylated regions were calculated using a 3 kb sliding window across the genome with a 600 bp step size. Overlapping regions were merged for each context separately.

BEDTools v2.24.0^52^ was used for all genomic features comparisons. Gencode v19 was used for genic annotations and transposable element coordinates were taken from RepeatMasker in the UCSC Genome Browser. HOMER{Heinz,2010} was used to identify Transcription factor binding motifs in regions of interest. Transcription factor binding sites were obtained from the ENCODE data track on UCSC Genome browser (http://genome.ucsc.edu/cgi-bin/hgTrackUi?db=hg19&=wgEncodeRegTfbsClusteredV3). LINE1 elements at non-CG hypermethylated differentially methylated regions were put into GREAT tool to predict gene targets.

## AUTHOR CONTRIBUTIONS

B.Y. conceived and designed the study, obtained funding, collected samples from collaborators, conducted all experiments, contributed to data analysis and wrote manuscript. N.D.J performed data analysis. S.L.B. contributed to data analysis and wrote manuscript. T.H.S., S.E.Z., and J.M. contributed to data analysis. T.S. and J.C. collected oocytes. R.D.H. contributed to data analysis and edited the manuscript.

## ACKNOWLEDGEMENTS

The authors would like to acknowledge Shahina Maqbool, PhD at Epigenomics and Genomics Center of Albert Einstein College of Medicine for her technical guidance on bisulfite sequencing.

## ACCESSION NUMBERS

The NCBI SRA accession number for the RNA-seq profiles of single oocytes reported in this paper is PRJNA508772. The link for reviewers is ftp://ftp-trace.ncbi.nlm.nih.gov/sra/review/SRP172762_20181207_112442_27e795eb0f314edf0479737480ab0f2a

## SUPPLEMENTAL INFORMATION

Three supplemental figures and three supplemental tables.

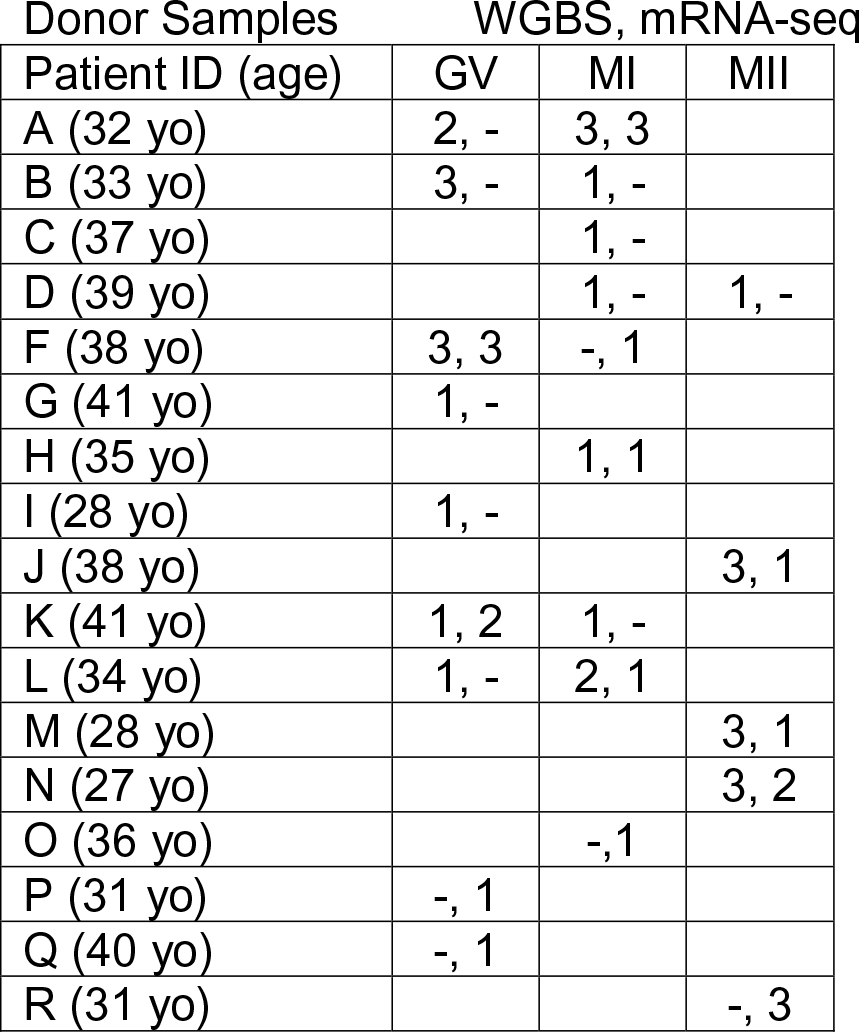
Donor Information, Related to Figure 1. The 17 donors from which single cell whole genome bisulfite sequencing (WGBS) or single cell mRNA-seq data was generated. The number of oocytes used for each method is written as: number of oocytes used for WGBS, number of oocytes used for mRNA-seq. A “-“ indicates no oocytes were used for that method. A blank cell indicates no oocytes from that stage were used. There is no sample E.

| Donor Samples | WGBS, mRNA-seq |  |  |
| --- | --- | --- | --- |
| Patient ID (age) | GV | MI | MII |
| A (32 yo) | 2, - | 3, 3 |  |
| B (33 yo) | 3, - | 1, - |  |
| C (37 yo) |  | 1, - |  |
| D (39 yo) |  | 1, - | 1, - |
| F (38 yo) | 3, 3 | -, 1 |  |
| G (41 yo) | 1, - |  |  |
| H (35 yo) |  | 1, 1 |  |
| I (28 yo) | 1, - |  |  |
| J (38 yo) |  |  | 3, 1 |
| K (41 yo) | 1, 2 | 1, - |  |
| L (34 yo) | 1, - | 2, 1 |  |
| M (28 yo) |  |  | 3, 1 |
| N (27 yo) |  |  | 3, 2 |
| O (36 yo) |  | -, 1 |  |
| P (31 yo) | -, 1 |  |  |
| Q (40 yo) | -, 1 |  |  |
| R (31 yo) |  |  | -, 3 |

Differentially Methylated Region Stats, Related to Figure 2. Statistics on differentially methylated regions (DMRs).

| Context | Comparison | DMRs | Hyper | Hypo | % distal | % distal DMR DHS overlap | % TE overlap |
| --- | --- | --- | --- | --- | --- | --- | --- |
| CpG | MI v GV | 1,559 | 974 | 585 | 57% | 92% | 97% |
| CHG | MI v GV | 789 | 545 | 244 | 70% | 85% | 99% |
| CHH | MI v GV | 21,453 | 15,471 | 5,982 | 75% | 84% | 99% |
| CpG | MII v MI | 905 | 575 | 330 | 54% | 93% | 96% |
| CHG | MII v MI | 3,834 | 3,469 | 365 | 71% | 86% | 99% |
| CHH | MII v MI | 53,469 | 42,923 | 10,546 | 73% | 85% | 99% |

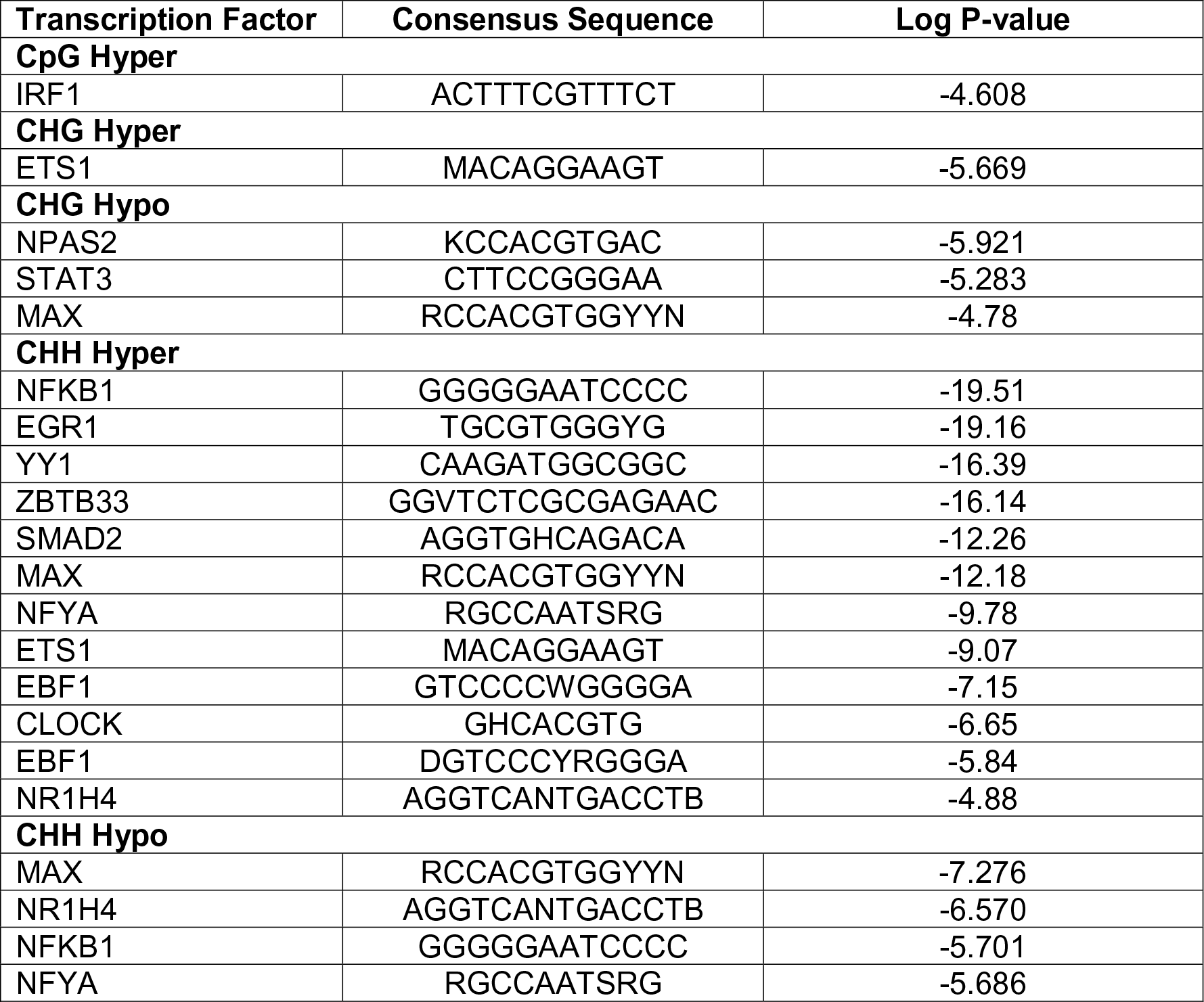
Expressed TFs with binding motifs in DMRs, Related to Figure 3. Transcription factor (TF) motifs found in MII-to-MI differential methylated regions (DMRs).

