## Supplemental figures for "Single-cell analysis of Non-CG methylation dynamics and gene expression in human oocyte maturation"

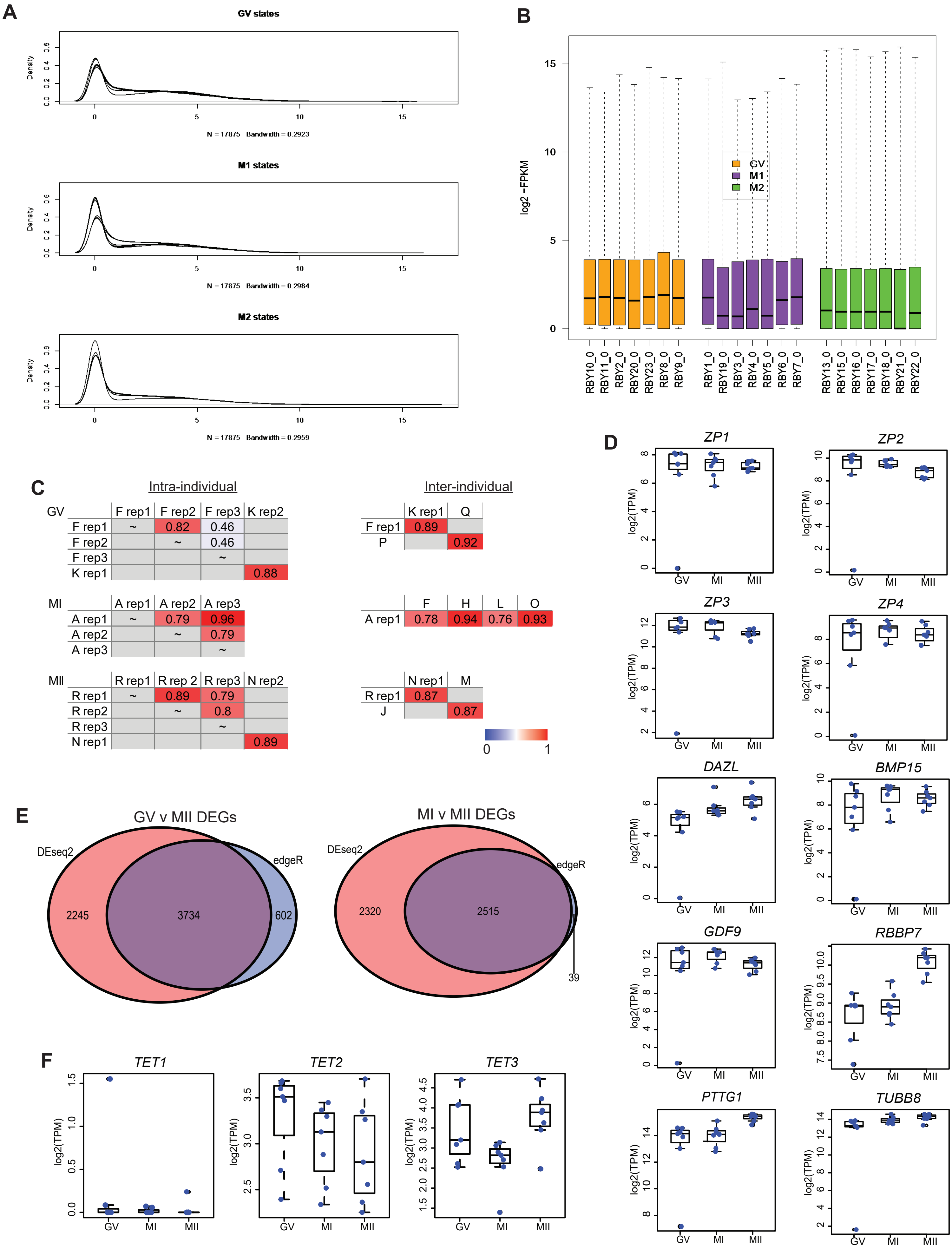


Supplemental Figure S1

Supplemental Figure S1
(A) Kernal density plot of the distribution of expression values for samples in each stage at 17.
(B) Box plots of log2(FPKM) for each individual sample.
(C) Heatmap of Pearson's correlations between samples from the same individual and bewteen different individuals. Not all comparisons shown.
(D) Box plots of select oocyte specific and MII upregulated genes. Blue dots are individual sample expression level.
(E) Venn diagram of the overlap between differentially expressed genes (DEGs) identified using DESeq2 or edgeR. DESeq2 DEGs were used for all downstream analysis.
(F) Expression levels of epigenetic regulators. Blue dots are individual sample expression level.

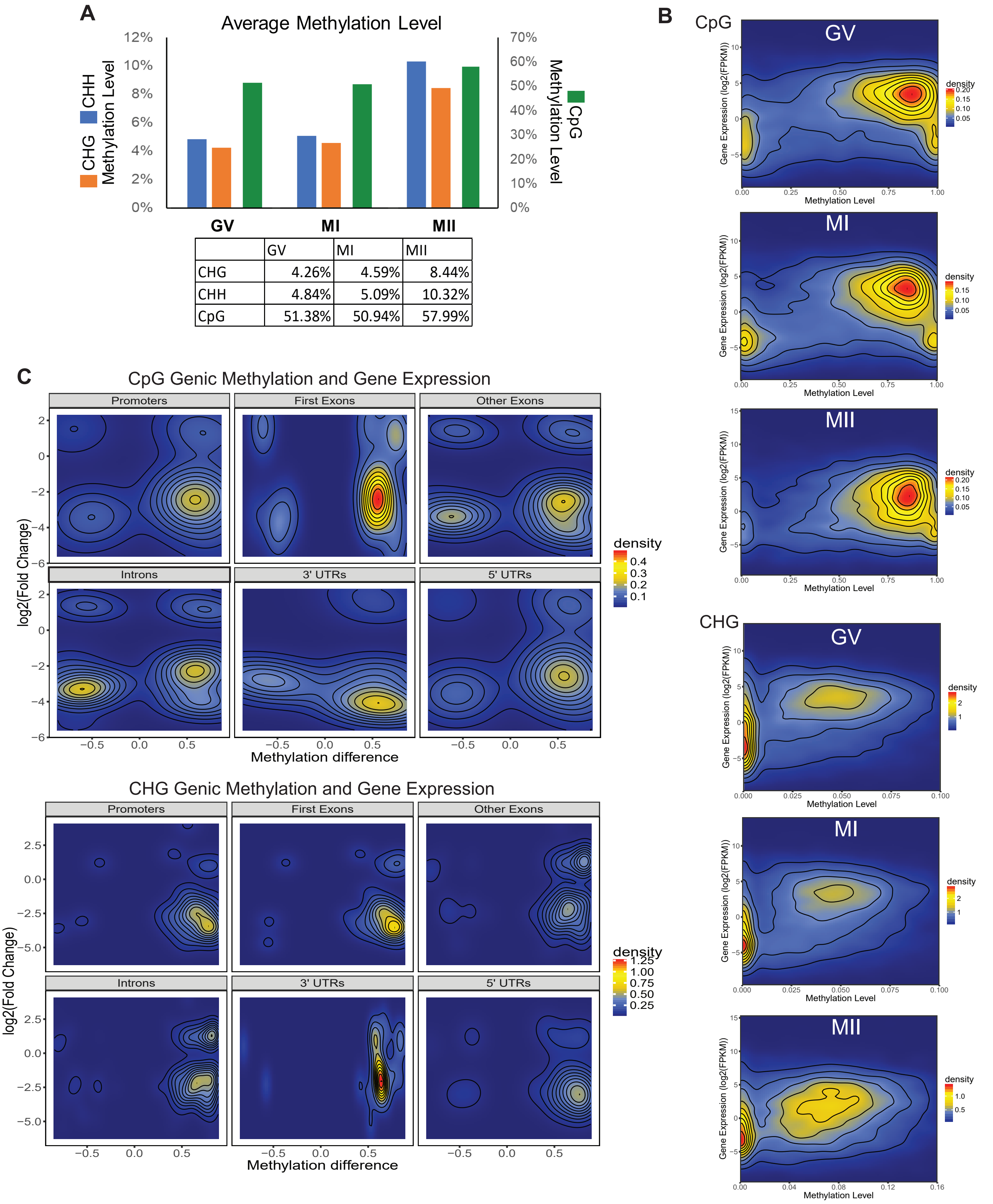


| | GV | MI | MII |
|---|---|---|---|
| CHG | 4.26% | 4.59% | 8.44% |
| CHH | 4.84% | 5.09% | 10.32% |
| CpG | 51.38% | 50.94% | 57.99% |

Supplemental Figure S2

(A) Average methylation level for each C context in each oocyte maturation stage. Adapted from Yu et al, 2017.

(B) Density plots of gene body methylation levels for CpG and CHG context and the corresponding gene's expression level in GV, MI and MII oocytes. Density color scale to the right of each respective plot.

(B) Density plots of genic methylation levels for CpG and CHG DMRs and the corresponding gene's fold change difference in MII-to-MI comparison. Density color scale to the right of each respective plot.

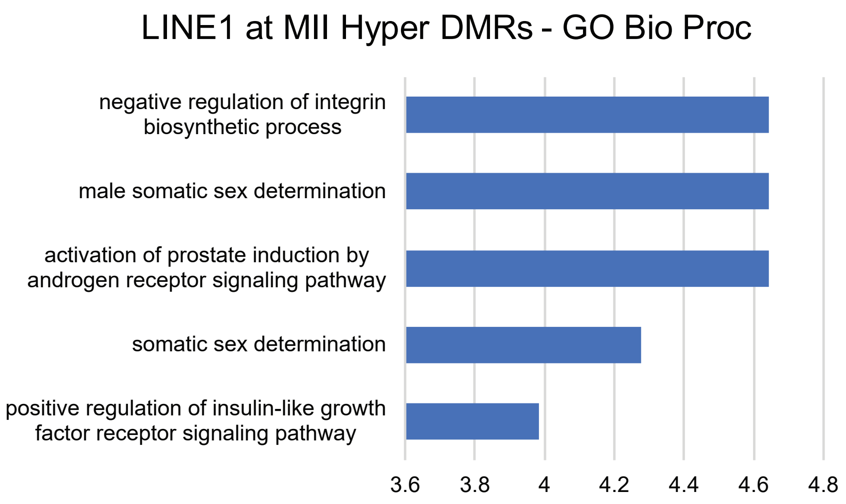


Supplemental Figure S3
GO Biological Processes for LINE1 elements at hyperDMRs in MII.
